# The pore-forming protein gasdermin D is a cellular redox sensor

**DOI:** 10.1101/2022.03.11.484021

**Authors:** Pascal Devant, Elvira Boršić, Elsy M. Ngwa, Jay R. Thiagarajah, Iva Hafner-Bratkovič, Charles L. Evavold, Jonathan C. Kagan

## Abstract

Reactive oxygen species (ROS) affect inflammation and immunity in a multitude of ways, in particular in the context of the signaling pathways that determine cell fate. Inflammasomes are multiprotein cytoplasmic complexes whose pyroptosis-inducing activities are controlled by ROS. Our knowledge of how ROS mediates inflammasome activities is largely based on studies of the initiating events in these pathways. Herein, we show that ROS controls the terminal events in the pyroptosis pathways. We found that ROS oxidizes the protein gasdermin D (GSDMD) and promotes its assembly into a death-inducing pore forming complex. Mechanistically, ROS enhances GSDMD-mediated pyroptosis in an intrinsic manner, likely through oxidative modification of a specific cysteine residue (C192). Diverse ROS sources promote GSDMD oxidation, ranging from homeostatic control via the Ragulator-Rag complex, to inducible control via diverse microbial products and environmental toxins. These findings expand the steps in the inflammasome pathway that are controlled by ROS and suggest that GSDMD operates as a pyroptosis-inducing redox sensor.

## Introduction

Reactive oxygen species (ROS) are produced by multiple mechanisms within cells, ranging from enzymatic activities to byproducts of aerobic respiration (Zorov et al., 2014). ROS can covalently modify proteins, DNA, and other biological molecules. The unregulated production of high amounts of ROS can be destructive to cells, as oxidized membranes may lose barrier function, oxidized DNA can result in mutation, and oxidized proteins can display altered biochemical activities. ROS species include unstable short-lived molecules such as superoxide and hydroxyl free radicals as well as more stable forms such as hydrogen peroxide. In extreme situations, such as following ionizing radiation, ROS can alter cellular behaviors in such catastrophic ways that lysis occurs (Juan et al., 2021). However, ROS species also serve regulatory functions through modifications of proteins, DNA, or membranes that alter signaling activities and cellular behaviors (Sies and Jones, 2020). Several examples of the influence of redox signaling have emerged, in particular in the context of environmental and inflammatory responses (Miller et al., 2010; Stottmeier and Dick, 2016).

Among the inflammatory mediators controlled by ROS is the interleukin-1 (IL-1) family of cytokines. Most members of this family do not contain an N-terminal secretion sequence (Dinarello, 2009). Consequently, IL-1β, the best-characterized member of this family, resides in the cytosol and is not released into the extracellular space via the conventional vesicle-mediated secretory pathway. Rather, IL-1β is commonly released after a lytic form of cell death known as pyroptosis, which is regulated by the protein gasdermin D (GSDMD) (Kayagaki et al., 2015; Shi et al., 2015; Afonina et al., 2015; Evavold et al., 2018; Heilig et al., 2018). GSDMD is synthesized as an inactive pro-protein that must be cleaved in order to self-assemble into an oligomeric complex that forms pores in the plasma membrane that can lead to cytolysis (*i*.*e*. pyroptosis) (Kayagaki et al., 2015; Shi et al., 2015; Afonina et al., 2015; Liu et al., 2016). GSDMD cleavage events are best-recognized to be mediated by inflammatory caspases, in particular those present in a large multiprotein complex called an inflammasome (Schroder and Tschopp, 2010). Inflammasomes are members of a growing family of innate immune supramolecular organizing centers (SMOCs), which are not present within resting cells but are assembled upon cellular encounters with infectious or non-infectious threats to the host (Kagan et al., 2014). The mechanisms by which inflammasomes are assembled have been the subject of intense investigation, with ROS being one of the first identified regulators of this process. Tschopp and colleagues recognized this link when they found that mitochondrial poisons that induce ROS via electron transport chain (ETC) disruption can promote the assembly of inflammasomes containing the protein NLRP3 (Zhou et al., 2011). In addition, more recent work has illustrated that microbial detection by Toll-like Receptors (TLRs) can lead to ROS production from mitochondria, which appears to position NLRP3 and caspase-1 on these organelles where they are poised to assemble into inflammasomes (West et al., 2011; Iyer et al., 2013; Elliott et al., 2018). ROS induced by TLR signaling also amplifies the expression of the genes encoding NLRP3 and IL-1β (Bauernfeind et al., 2011; van de Veerdonk et al., 2010). Based on these studies, the prevailing concept is that ROS acts at the apex of inflammasome pathways to regulate their function. Whether and how ROS acts at other stages of pyroptosis pathways remains undefined.

In a genome-wide screen to identify genes that act downstream of inflammasomes and GSDMD cleavage, we recently identified the Ragulator-Rag complex, which is best-known for its control of cellular metabolism (Evavold et al., 2021). We found that macrophages lacking the Ragulator-Rag components RagA or RagC displayed defects in GSDMD pore formation and pyroptosis. A link to ROS regulation was suggested, as defects in GSDMD pore forming activity in RagA- and RagC-deficient cells could be rescued by the supply of exogenous ROS or induction of endogenous ROS by mitochondrial poisons. A separate study also reported a role for Ragulator-Rag in pyroptosis (Zheng et al., 2021). While both studies utilized genome-wide genetic screens to identify regulators of GSDMD-mediated pyroptosis, the experimental systems were quite different. The latter report studied the pyroptosis pathway that is induced when TLRs are activated in the presence of chemical inhibitors of the kinase TAK1. Under these conditions, the kinase RIPK1 activated caspase-8, leading to GSDMD-dependent pyroptosis. In contrast, our study utilized a genetic system whereby macrophages were engineered to express the pore forming domain of GSDMD (known as NT-GSDMD) under the control of a doxycycline (Dox)-inducible promoter. Thus, whereas our study bypassed upstream regulatory events that lead to GSDMD cleavage, the study by Zheng and colleagues stimulated proteins that act upstream of GSDMD.

Herein, we sought to better define the role of Ragulator-Rag and ROS regulation in GSDMD activities. We used our Dox-inducible NT-GSDMD system to focus our studies on the terminal events in pyroptosis. Moreover, to further enable our studies, we utilized a version of GSDMD that contains an I105N mutation. As described initially by Dixit and colleagues, this NT-GSDMD I105N retains the ability to form membrane pores, but does so more slowly than its wild type counterpart (Kayagaki et al., 2015; Aglietti et al., 2016). This variant allowed for an increased window of time to study GSDMD through microscopic and biochemical analysis prior to cell rupture. Our studies revealed that under steady state conditions, Ragulator-Rag is required for basal ROS production that enables NT-GSDMD pore formation and pyroptosis. Diverse innate immune signaling pathways or environmental perturbations that induce ROS can bypass the requirement of Ragulator-Rag and promote NT-GSDMD oligomerization and pore formation. Finally, we found that GSDMD is directly oxidized at multiple cysteine residues, with C192 being required for ROS-mediated potentiation of GSDMD pore formation and pyroptosis. These findings reveal GSDMD as a sensor of cytosolic redox states whose oxidation determines cell fate.

## Results and Discussion

The two studies characterizing Ragulator-Rag in pyroptosis characterized distinct subunits of this protein complex (Evavold et al., 2021; Zheng et al., 2021). To compare the findings that were made, we created a common set of genetic tools and performed assays using pyroptosis stimuli that had been previously used in each of these studies. We generated immortalized murine bone marrow-derived macrophages (iBMDMs) deficient for the Ragulator-Rag component RagA, the central GTPase that controls Ragulator functions. We electroporated Cas9-expressing iBMDMs with three different sgRNAs targeting *Rraga*, the gene encoding RagA. Complete ablation of the RagA protein was observed by immunoblot analysis in the resulting clonal iBMDM lines (Figure 1A). These cells were co-stimulated with the TLR4 ligand bacterial lipopolysaccharide (LPS) and the chemical 5z7, an inhibitor of TAK1. This treatment has been used to stimulate RIPK1- and caspase-8-mediated pyroptosis (Orning et al., 2018; Zheng et al., 2021). Pyroptosis was assessed by monitoring the release of the cytosolic enzyme lactate dehydrogenase (LDH) into the extracellular space. Additionally, membrane permeabilization was assessed by staining cells with propidium iodide (PI), a membrane impermeable dye that fluoresces upon binding intracellular nucleic acids. Cells that received a non-target control sgRNA (referred to as WT) stained strongly for PI and released LDH after LPS+5z7 treatment (Figure 1B, C). In contrast, RagA- deficient cells were protected from pyroptosis induction using both of these assays (Figure 1B, C). Thus, similar to studies of other components of the Ragulator-Rag complex (Zheng et al., 2021), RagA is required for the pyroptosis induced by LPS+5z7.

**Figure 1:**
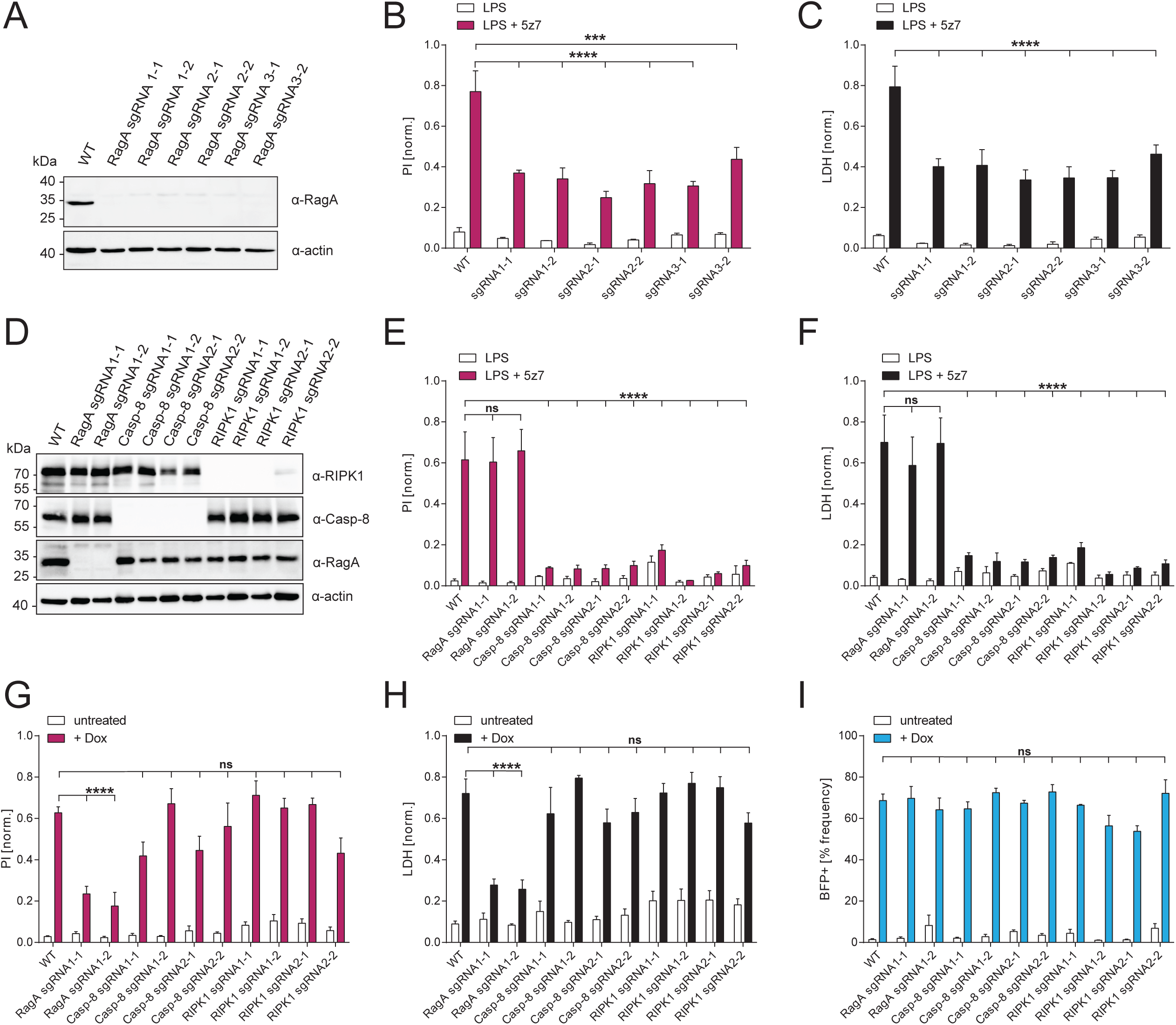
Dox-induced expression of NT-GSDMD leads to RIPK1- and caspase-8-independent pyroptosis. (A) Immunoblot analysis of whole-cell lysates of iBMDM clonal cell lines in which expression of RagA was disrupted by CRISPR/Cas9. (B, C) RagA-deficient iBMDM or non-target sgRNA (WT) control cells were either treated with LPS (1 μg/ml) alone or co-treated with LPS (1 μg/ml) and the TAK1 inhibitor 5z7 (250 nM) for 3 h. PI uptake and LDH release into cell culture supernatants were quantified to assess plasma membrane disruption and cellular lysis, respectively. (D) Immunoblot analysis of whole-cell lysates of iBMDMs expressing NT-GSDMD under Dox-inducible promoter in which expression of RagA, caspase-8, or RIPK1 was disrupted by CRISPR/Cas9. (E, F) iBMDMs expressing NT-GSDMD under Dox-inducible promoter in which expression of RagA, caspase-8, or RIPK1 was disrupted by CRISPR/Cas9 were treated with LPS (1 μg/ml) alone or co-treated with LPS (1 μg/ml) and the TAK1 inhibitor 5z7 (250 nM) for 6 h. PI uptake and LDH release into cell culture supernatants were quantified to assess plasma membrane disruption and cellular lysis, respectively. (G, H, I) iBMDMs expressing NT-GSDMD under Dox-inducible promoter in which expression of RagA, caspase-8, or RIPK1 was disrupted by CRISPR/Cas9 were treated with Dox (0.5 μg/ml) for 16 h. PI uptake and LDH release into cell culture supernatants were quantified to assess plasma membrane disruption and cellular lysis, respectively. Fraction of BFP-positive cells was measured by flow cytometry. Cells indicated as WT were electroporated with a non-targeting sgRNA. Data are represented as mean ± SEM of three independent experiments. Immunoblots show representative result of three independent repeats. Statistical significance was determined by two-way ANOVA: *p < 0.05; **p < 0.01; ***p < 0.001; ****p < 0.0001.

We considered these findings in the context of our recent report that RagA was required for pyroptosis induced by the Dox-mediated expression of NT-GSDMD (Evavold et al., 2021). It was possible that Dox treatments may stimulate RIPK1 and caspase-8, leading to NT-GSDMD mediated pore formation. If this possibility was correct, then impairing the function of RIPK1 or caspase-8 should lead to decreased pyroptosis upon Dox-induction of NT-GSDMD. We first tested this hypothesis using a small molecule inhibitor-based approach. We Dox-induced NT-GSDMD in the presence of chemical inhibitors of RIPK1 (7-Cl-O-Nec-1), the related kinase RIPK3 (GSK’872) or the pan-caspase inhibitor zVAD. None of these inhibitors interfered with NT-GSDMD induced cell death, as indicated by LDH release (Supplementary Figure 1A). The inhibitor concentrations used in this analysis were functional, as 7-CL- O-Nec-1 inhibited LPS+5z7-induced cell death (Supplementary Figure 1B, C). Additionally, ZVAD treatments converted LPS+5z7 induced cell death into one that is RIPK3-dependent (CSK’872 sensitive), and ZVAD treatments blocked all evidence of GSDMD cleavage (Supplementary Figure 1B-D).

To corroborate these findings, we generated cells lacking the genes encoding caspase-8 or RIPK1 using CRISPR-Cas9 (Figure 1D). We functionally confirmed RIPK1 and caspase-8 deficiencies, as the cells lacking these proteins were protected from LPS+5z7-induced pyroptosis, even after prolonged treatment of up to 6 hours (Figure 1E, F). Interestingly, at this late time point, RagA-deficient cells died to a similar extent as WT control cells (Figure 1E, F). These results indicate that RIPK1 and caspase-8 are required for LPS+5z7 induced cell death at all time points examined, whereas RagA is most important for the rapid pyroptotic response induced by LPS+5z7. Similar to the findings made with chemical inhibitors, neither caspase-8 nor RIPK1 deficiencies prevented pyroptosis induced by NT-GSDMD (Figure 1G, H). In contrast, RagA-deficient cells were protected from cell death under the same conditions (Figure 1G, H). Using the BFP tag that is appended to the C-terminus of NT-GSDMD, we verified that the frequency of BFP-positive cells was unchanged under all experimental conditions (Figure 1I). These collective findings indicate that RagA is required for pyroptosis induced by Dox-mediated NT-GSDMD, as well as early pyroptotic responses induced by LPS+5z7 treatments.

The common node in the pyroptotic events induced by LPS+5z7 and Dox-treatments is GSDMD, suggesting that RagA acts at this stage in the pyroptosis pathway. To address the functions of GSDMD specifically, our subsequent work utilized the Dox-inducible system for NT-GSDMD activities. To better understand the link between RagA and GSDMD, we focused on the aforementioned role of Ragulator-Rag in ROS regulation. We assembled a panel of agents that increase cellular ROS in distinct manners. This panel consists of two microbial products, LPS and fungal β-glucans, the environmental toxin sodium arsenite, and two mitochondrial ETC inhibitors, rotenone and TTFA (Hsu and Wen, 2002; Hei et al., 1998; Underhill et al., 2005). Rotenone is a ROS fluxing agent whereas treatment with TTFA is not expected to increase cellular ROS (Zhou et al., 2011). Live cell confocal microscopy was used to assess ROS production by staining with the redox-sensitive fluorescent probe CM-H_2_DCFDA.

Using this single cell analysis, we found cellular ROS levels were lower in RagA-deficient cells than in empty vector (WT) control cells (Figure 2A, B) consistent with orthogonal ROS measurements under basal and Dox-induced conditions described previously (Evavold et al., 2021). Treatment with Dox to induce NT-GSDMD expression led to an increase in cellular ROS in WT cells, but not in RagA-deficient cells (Figure 2A, B). Interestingly, the ROS production defect in RagA-deficient cells was bypassed by treatments with LPS, β-glucan, sodium arsenite or rotenone (Figure 2A, B). TTFA did not induce any ROS in these cells (Figure 2A, B). Based on the ability of these agents to bypass the ROS production defects of RagA-deficient cells, we examined their ability to rescue defects in NT-GSDMD oligomer formation, PI staining and LDH release. Dox-induction of NT-GSDMD led to an increase in PI staining and LDH release in WT cells, but not RagA-deficient cells (Figure 2C, D). We found that all the agents that rescued the ROS production defects in RagA-deficient cells (LPS, β-glucan, sodium arsenite, rotenone) also rescued PI staining and LDH release defects in these cells (Figure 2C, D). Importantly, the rescue of PI staining and LDH release was accompanied by an increase in GSDMD oligomerization, as determined by immunoblot under non-reducing conditions (Figure 2E) (Liu et al., 2016). These data therefore identify a diverse array of ROS-fluxing stimuli, including those that are commonly used to prime cells for inflammasome responsiveness, as capable of restoring GSDMD pore formation and pyroptosis in RagA-deficient cells.

**Figure 2:**
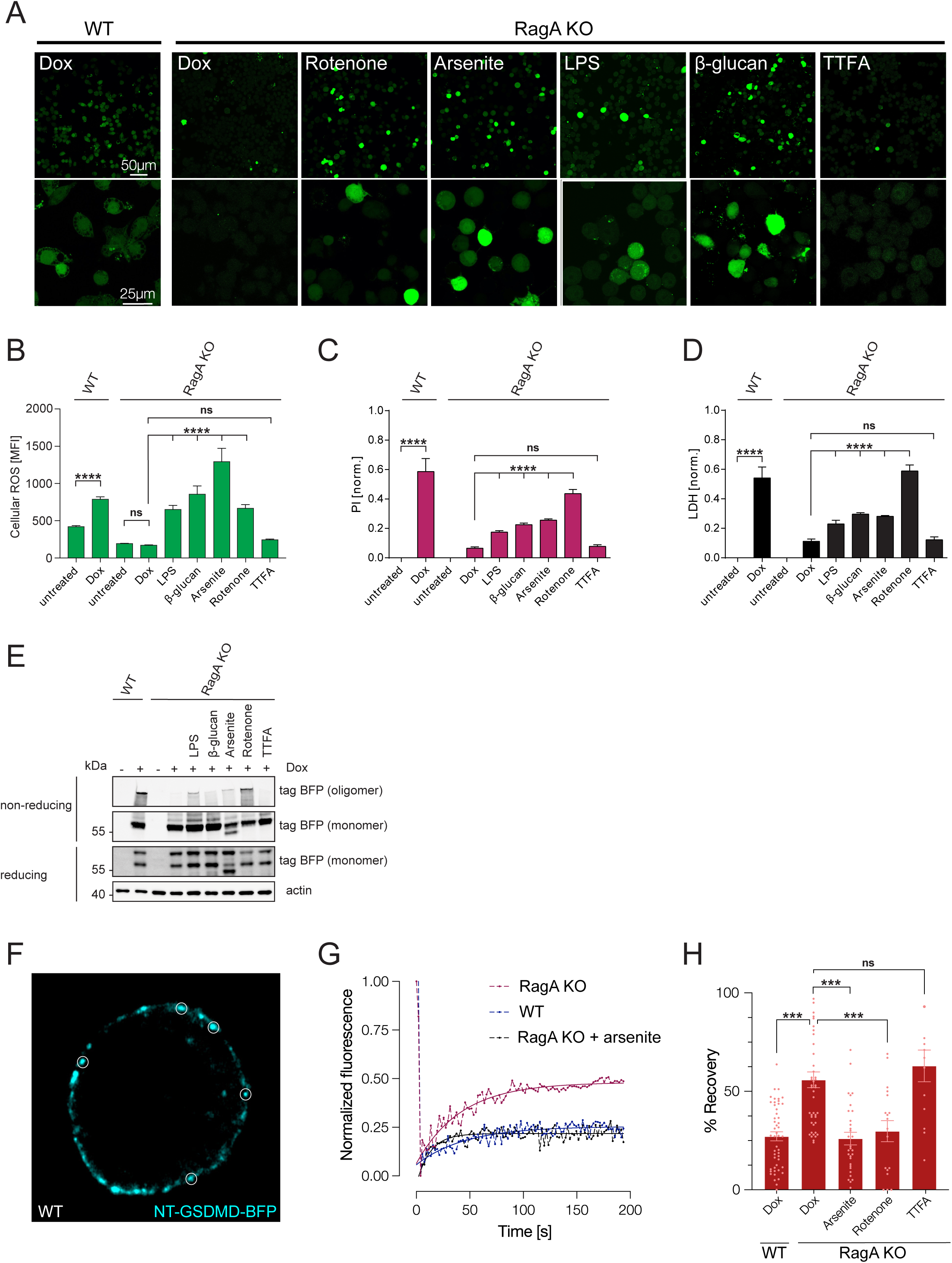
Priming with microbial products or environmental toxins induces ROS and promotes GSDMD pore formation. RagA-deficient or WT control iBMDMs expressing NT-GSDMD fusion protein under Dox-inducible promoter were treated with Dox (2 μg/ml in A, B, E or 0.5 μg/ml in C, D) for 8 h. Indicated PAMPs (1 μg/ml of LPS; 1 mg/ml of β-glucan) were present throughout entire stimulation period, while toxins (20 μM sodium arsenite; 5 μM rotenone; 100 μM TTFA) were added after 4 h. (A, B) Cells were stained using the ROS-reactive dye CM-H2DCFDA and signal was quantified by fluorescence microscopy. (C) PI uptake was quantified to assess plasma membrane disruption. (D)LDH release into cell culture supernatants was quantified as a measure of cellular lysis. (E)GSDMD oligomerization was assessed by non-reducing SDS-PAGE followed by immunoblot. (F)Representative confocal image of WT control cell after Dox-induction of NT-GSDMD-BFP for 8 h showing membrane distribution of NT-GSDMD-BFP. Circles indicate areas for measurement of fluorescence recovery after photobleaching (FRAP). (G)Representative FRAP curves from WT control cells, RagA KO and RagA KO cells co-treated with sodium arsenite. (H)Mean proportion of fluorescence recovery following photobleaching of BFP signal from NT-GSDMD-BFP in WT and RagA KO macrophages after Dox induction (2 μg/mL) for 8 h. Arsenite (20 μM), Rotenone (5 μM) or TTFA (100 μM) was added after 4 h. Cells indicated as WT were transduced with an empty vector construct. Data are represented as mean ± SEM of three independent experiments. Immunoblots and microscopy images show representative result of three independent repeats. Statistical significance was determined by one-way ANOVA (B-D) or two-way ANOVA with Tukey’s post-hoc testing (H): *p < 0.05; **p < 0.01; ***p < 0.001; ****p < 0.0001.

To complement the biochemical analysis of ROS-induced GSDMD oligomer formation, we examined oligomerization of GSDMD within individual cells *in situ*. We carried out fluorescence recovery after photobleaching (FRAP) experiments on BFP-tagged NT-GSDMD. Following induction of NT-GSDMD in RagA-deficient or WT control cells, we targeted membrane localized BFP fluorescence areas for photobleaching and monitored subsequent fluorescence recovery (Figure 2F). Multimerization of a membrane-bound protein into a larger complex leads to reduced two-dimensional diffusion within the membrane, as compared to monomers. This reduced diffusion of a large complex leads to longer fluorescence recovery times following photobleaching, as well as a reduction in the total mobile fraction or percent recovery over the timescale of the experiment (Haggie and Verkman, 2008). Consistent with this notion, RagA-deficient cells (where GSDMD oligomerization is impaired) exhibited an increased mobile fraction compared to WT control cells (Figure 2G, H). Consistent with our biochemical analysis for NT-GSDMD oligomerization, treatment with the ROS-inducing agents rotenone or sodium arsenite reduced the percent recovery in RagA-deficient cells (Figure 2G, H). In contrast, TTFA treatments, which do not induce ROS, did not alter fluorescence recovery of NT-GSDMD in RagA-deficient cells (Figure 2G, H). These experiments in intact live macrophages support the conclusion that ROS controls GSDMD oligomerization and pore formation.

Redox-regulation of proteins may occur via direct modification of thiol-containing amino acid residues such as cysteines. A possible mechanism for ROS-mediated GSDMD pore regulation would therefore be direct oxidation of cysteine residues within NT-GSDMD. If this hypothesis is correct, mutation of cysteine residues to a non-oxidizable amino acid (*e*.*g*. alanine) should affect the ability of NT-GSDMD to form pores and induce cell death. Several cysteines in the N-terminal domain of GSDMD, including C191/192 and C38/C39 in human/murine GSDMD, have been implicated in GSDMD oligomerization and pore formation (Liu et al., 2016; Rathkey et al., 2018; Humphries et al., 2020). To investigate the importance of these cysteine residues for GSDMD-mediated cell death, we engineered GSDMD-deficient iBMDMs to express NT-GSDMD carrying C39A or C192A mutations, or a NT-GSDMD variant where all six cysteines have been mutated to alanine (No Cys), under the control of a Dox-inducible promoter. Dox treatments of iBMDMs led to the production of comparable amounts of all NT-GSDMD proteins, but none of the mutant NT-GSDMD variants exhibited a decreased ability to induce pyroptosis (Figure 3A-C). This finding was surprising, as previous studies using transient transfections of HEK293 cells concluded that GSDMD lacking C192 is deficient in pore forming abilities (Rathkey et al., 2018; Liu et al., 2016; Humphries et al., 2020; Hu et al., 2020). Based on the suggested role for ROS in GSDMD pore formation, we wondered whether an additional supply of ROS might uncover a phenotype in the context of cysteine substitution. To examine this possibility, we induced NT-GSDMD variants with Dox and provided exogenous ROS by co-treating cells with H_2_O_2_. Under these conditions, we observed an enhancement of pore forming activities induced by WT NT-GSDMD and the C39A mutant (Figure 3B, C). In contrast, no ROS-mediated enhancement of PI staining or LDH release was observed in cells expressing the C192A or the No Cys mutant (Figure 3B, C). These findings support the role of ROS as a potentiator of GSDMD pore forming activities and establish C192 as a specific regulator of these events.

**Figure 3:**
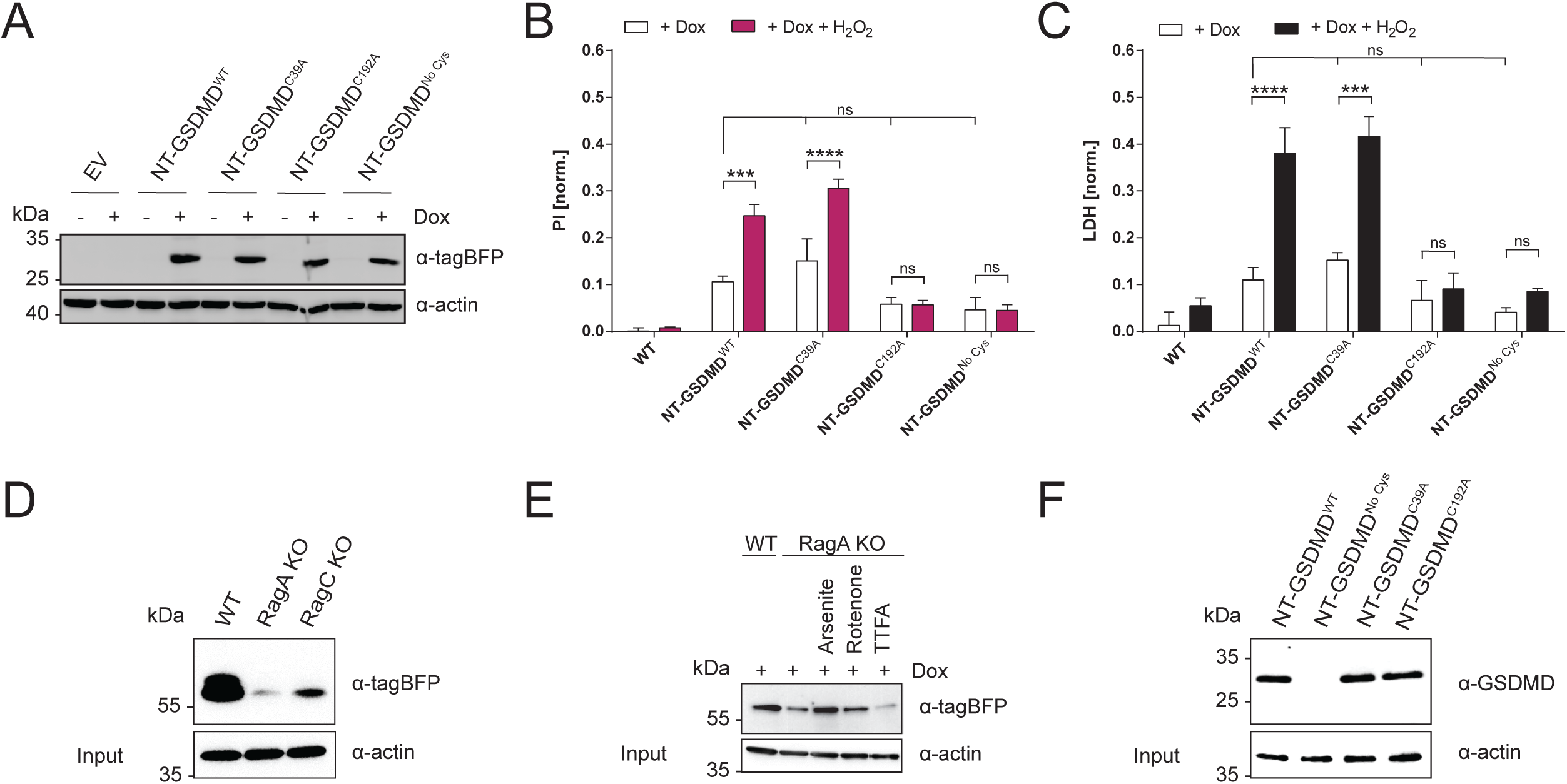
ROS oxidizes cysteines on NT-GSDMD, leading to maximal pyroptosis activities. (A)Immunoblot analysis of whole-cell lysates of GSDMD KO iBMDM expressing indicated NT-GSDMD mutants under a Dox-inducible promoter or tranduced with an empty vector (EV). Cells were treated with 1 μg/ml of Dox for 4 h or left untreated. (B, C) GSDMD KO iBMDMs expressing indicating NT-GSDMD variants under a Dox-inducible promoter or transduced with an empty vector (EV) were treated with Dox (1 μg/ml) for 8 h. H_2_O_2_ was added after 6 h to a final concentration of 625 μM. PI uptake was quantified to assess plasma membrane disruption and LDH release into cell culture supernatants was quantified as a measure of cellular lysis. (D, E) Pulldown and immunoblot analyses to assess relative cysteine oxidation in RagA-deficient, RagC-deficient or empty-vector treated iBMDMs (WT) expressing NT-GSDMD-BFP fusion protein under Dox-inducible promoter. Cells were treated with Dox (2 μg/ml) for 8 h. Indicated toxins (20 μM sodium arsenite; 5 μM rotenone; 100 μM TTFA) were added after 4 h. (I)Pulldown and immunoblot analyses to assess relative cysteine oxidation in GSDMD KO cells expressing indicated NT-GSDMD mutants under Dox-inducible promoter. Cells were treated with Dox (2 μg/ml) for 8 h. Data shown in A,B,C were generated using representative clonal cell populations, while data presented in F were generated using a bulk population of cells. Clonal populations used in A,B,C were directly derived from bulk populations used in F. Data are represented as mean ± SEM of three independent experiments, except for EV in B and C, which is mean ± SEM of two independent experiments. Immunoblots show representative result of three independent repeats. Statistical significance was determined by two-way ANOVA with Tukey’s post-hoc testing: *p < 0.05; **p < 0.01; ***p < 0.001; ****p < 0.0001.

We therefore investigated whether any cysteines in NT-GSDMD are oxidized by ROS. We induced NT-GSDMD and analyzed relative cysteine oxidation by pulling down oxidized cysteine (sulfenic acid) residues using a reactive biotin-maleimide probe (Saurin et al., 2004). We detected a signal for GSDMD in WT control cells upon Dox-treatment (Figure 3D), indicating that at least one cysteine in NT-GSDMD is oxidized. This signal of NT-GSDMD oxidation was greatly diminished in RagA- or RagC- deficient cells (Figure 3D), a finding consistent with lower ROS in the absence of Ragulator-Rag functions (Figure 2A, B). However, when RagA-deficient cells were co-treated with the ROS-inducing agents rotenone or sodium arsenite, NT-GSDMD oxidation was partially restored (Figure 3E). To determine whether cysteine oxidation within NT-GSDMD is site-specific, we determined the relative oxidation of select NT-GSDMD mutants (C192A, C39A and No Cys mutant). We observed no notable reduction in protein oxidation when examining the C39A or C192A mutants, but complete abrogation of ROS- modifications in the No Cys mutant, which lacks all major oxidizable amino acids (Figure 3F). The simplest interpretation of this data is that multiple cysteine residues within NT-GSDMD can be oxidized by ROS, but only oxidation at site C192 affects its pore-forming abilities that lead to pyroptosis.

Our collective data demonstrates a link between ROS regulation and GSDMD activities. ROS has been long-recognized as an inducer of inflammasome assembly via NLRP3 (Zhou et al., 2011), and has recently been implicated in the regulation of the inflammasome adaptor protein ASC (Li et al., 2021). Importantly, both of these ROS-controlled regulatory mechanisms of inflammasome activity operate upstream of GSDMD cleavage. A central point of novelty that emerges from this study is that ROS can directly potentiate NT-GSDMD oligomerization, independently of upstream inflammasome effects. Based on the established role of ROS at the apex of the NLRP3 pathway, we propose the idea of a ROS regulatory cascade, akin to phosphorylation cascades, where multiple nodes in a pathway are subject to the same type of regulation (ROS or phosphorylation). This idea has important implications for the use of LPS and other innate immune agonists that flux ROS as priming agents to promote inflammasome activities.

We propose an additional role for the priming step in inflammasome activation (often referred to as signal 1). In addition to the invoked transcriptional changes (upregulation of inflammasome components and pro-IL-1β), priming agents also induce ROS that may potentiate GSDMD oligomer formation and pyroptosis. Moreover, the common association of ROS with mitochondrial dysfunction and microbial infection suggests that ROS may act as a secondary messenger within host defense pathways to communicate threat level through regulation of distinct inflammasome-related proteins.

Mechanistically, our finding that cysteine residues in NT-GSDMD are oxidized by ROS in a Rag- Ragulator-dependent manner is notable, as cysteine residues are known targets of several metabolite-based and small-molecule GSDMD inhibitors. For example, C192 is targeted by the cysteine-reactive inhibitors disulfiram, necrosulfamide and dimethyl fumarate, highlighting that this residue is accessible for posttranslational modifications (PTMs)(Hu et al., 2020; Rathkey et al., 2018; Humphries et al., 2020). However, our functional analysis of Dox-inducible NT-GSDMD lacking select cysteine or all cysteine residues did not yield a strong defect in pyroptosis induction under all conditions examined. Only upon exogenous ROS exposure, which potentiated WT GSDMD pore forming activities, was a role for C192 revealed as critical for maximal pyroptosis induction. The precise mechanism of how oxidation of cysteines within GSDMD contribute to pore formation remains to be determined. ROS-mediated oxidation of cysteine residues could potentially aid the addition or removal of an activating or inhibitory PTM, respectively. Our data is consistent with a recent report that suggests that host metabolic state may modify GSDMD with an inhibitory mark (Humphries et al., 2020; Bambouskova et al., 2021). As ROS can enhance pyroptosis in a C192-dependent manner, our data may represent removal of an inhibitory mark in a subset of GSDMD molecules. Furthermore, it remains to be determined what role cysteine residues within GSDMD, and their oxidation, play after natural pathway stimulation. Overall, we propose that GSDMD may be considered a pyroptosis-inducing sensor of cellular redox states, a finding that provides a mandate to explore GSDMD cysteine oxidation as a biomarker of productive inflammasome activation.

## Materials and Methods

### Reagents and antibodies

*E. coli* LPS (serotype O:111 B4) was purchased from Enzo Biosciences as a ready-to-use stock solution of 1 mg/ml. β-glucan-peptide was from Invivogen and dissolved at 5 mg/ml in sterile PBS. zVAD-FMK (Invivogen) was resuspended in sterile DMSO to a concentration of 20 mM and used at a final concentration of 20 μM or 50 μM. Propidium iodide solution (1 mg/ml) was purchased from MilliporeSigma. RIPK1 inhibitor (7-Cl-O-Nec-1) and RIPK3 inhibitor (GSK’872) were both from MilliporeSigma. TAK1 inhibitor (5z7) was purchased from Sigma and resuspended in sterile DMSO at 2.5 mM. Doxycycline hyclate was from Sigma and resupended at 1 mM in sterile H_2_O. Sodium arsenite (MilliporeSigma) was dissolved in H_2_O to a concentration of 100 mM and sterile-filtered through a 0.2 μm syringe filter before use. Recombinant murine LBP was purchased from R&D, diluted to 1 mg/ml in sterile PBS and snap-frozen in LN2. G418 was from Invivogen and puromycin was from Gibco. Stabilized hydrogen perodxide (H_2_O_2_) solution (30 %) was purchased from Sigma-Aldrich.

Following antibodies were used as primary antibodies for immunoblotting: mouse monoclonal anti-β-actin (MilliporeSigma, 1:5000), rabbit monoclonal anti-GSDMD (Abcam, 1:1000), rabbit monoclonal anti-RagA (Cell Signaling, 1:1000), rabbit monoclonal anti-caspase-8 (Cell Signaling, 1:1000), rabbit monoclonal anti-RIPK1 (Cell signaling, 1:1000), rabbit polyclonal anti-tagRFP (Evrogen, 1:1000). HRP-conjugated goat anti-mouse and goat anti-rabbit IgG (H+L) secondary antibodies were from Jackson ImmunoResearch and diluted 1:5000.

### Plasmid Constructs

Gene encoding mouse full-length GSDMD was ordered from Synobiological. gBlock encoding full-length mouse GSDMD with all cysteines in the N-terminal domain mutated to alanine (No Cys mutant) was ordered from Integrated DNA Technologies. N-terminal GSDMD (1-276) was amplified by PCR and subcloned into pRETROX TRE3G vector using BamHI and EcoRI restriction sites. I105N, C39A or C192A point mutations were introduced by site-directed mutagenesis. All constructs were verified by Sanger sequencing.

### Cell lines

All cells were cultured in humidified incubators at 37 °C and 5% CO_2_. Immortalized bone marrow-derived macrophages (iBMDMs) and Platinum GP cells (Cell Biolabs) were cultured in DMEM supplemented with 10% fetal bovine serum (FBS), Penicillin + Streptomycin, L-Glutamine and Sodium pyruvate, herafter referred to as complete DMEM (cDMEM) and cultured in tissue-culture treated 10 cm dishes or T175 tissue culture flasks (Corning). iBMDMs and Platinum-GP cellswere passaged using PBS + 4 mM EDTA or 0.25% trypsin + EDTA (Gibco), respectively. iBMDMs derived from a Cas9 knock-in mouse or a GSDMD KO mouse and Cas9-expressing iBMDMs expressing a NT-GSDMD-tagBFP fusion protein with an I105N mutation (referred to as NT-GSDMD) under the control of a Dox-inducible promoter (parental line used to generate KOs for Figure 1; empty vector-transduced and RagA KO cells used in Figure 2, 3 and 4) were generated previously (Evavold et al., 2021).

Dox-inducible NT-GSDMD(I105N) variants were introduced into GSDMD KO iBMDMs by sequential transduction with two pantropic retroviruses. First, the TET3G tetracycline transactivator protein was introduced. To do so, 1.2 × 10^6^ Platinum-GP retrovirus packaging cells were seeded per well in a 6-well tissue culture plate. On the next day, cells were transfected with pCMV-VSV-G (1.5 μg) and pRETROX TET3G (2.5 μg) (Clontech) using Lipofectamine 2000 (10 μL). The same day 3 × 10^5^ GSDMD KO cells were seeded per well of 6-well plate. The next day, medium was replaced with 1 ml of complete medium with polybrene (2 μg/ml) and retroviral supernatant was filtered through 0.45-micron syringe filter and added to cells. Three days after transduction, transduced cells were selected by growth in cDMEM including 1.5 mg/ml G-418. Clonal cell lines were derived by limited serial dilution. GSDMD-NT(I105N) variants were introduced into single cell GSDMD-KO/Tet3G cell line by transduction of respective pantropic viruses using above described protocol where pRETROX TRE3G vectors encoding GSDMD-NT(I105N) variants were used as transfer plasmids. Selection of transduced cells was performed by growth in 10 μg/ml puromycin and 1.5 mg/ml G-418 medium.

### Generation of KO cell lines using CRISPR-Cas9 technology

CRISPR KO cell lines were generated by electroporation of synthetic sgRNAs into Cas9-expressing iBMDMs using the Neon transfection system (Thermo Fisher) as described before (Evavold et al., 2021). Briefly, 1.2×10^6^ Cas9-expressing iBMDMs were resuspended in 120 μl of R buffer, mixed with 2 μl of sgRNA (ordered from IDT and resuspended in nuclease-free water at a concentration of 100 μM) and electroporated using a 100 μl electroporation pipette tip with two 10 ms pulses at a voltage of 1400 V. Cells were then dispensed directly into a 6-well plate containing 3 ml of cDMEM and cultured for 3-5 days before assessing bulk KO efficiency by immunoblot. These cell lines were further single cell cloned by limited serial dilution to obtain clonal KO cell populations with complete ablation of the target protein. sgRNA sequences were previously published or pre-designed by IDT (Non-target sgRNA: AAAUGUGAGAUCAGAGUAAU; Caspase-8 sgRNA1: CTTCCTAGACTGCAACCGAG; Caspase-8 sgRNA2: GTGGGATGTAGTCCAAGCAC; RagA sgRNA1: GGTTCCCCAAGAATCGGACG; RagA sgRNA2: GATCAGCTGATAGACGATGC; RagA sgRNA3: GTGGGAGTGCTCCACGTCGA; RIPK1 sgRNA1: TACACGTCCGACTTCTCCGT; RIPK1 sgRNA2: AGTCGAGTGGTGAAGCTACT).

### GSDMD oligomerization assay

0.5×10^6^ iBMDMs expressing NT-GSDMD under a Dox-inducible promoter were seeded in 12-well plates in 1 ml of cDMEM and incubated at 37 °C and 5% CO_2_ overnight. Media was exchanged for 1 mL of Opti-MEM containing 2 μg/ml of Dox to induce the expression of GSDMD-NT-tagBFP for 8 h at 37 °C and 5% CO_2_. As indicated, cells were cotreated with ROS-inducing agents or PAMPs. 50 μl of Opti-MEM containing rotenone (5 μM final concentration), TTFA (100 μM final concentration) or sodium arsenite (20 μM final concentration) was added to the cell culture media after 4 h. LPS (1 μg/ml) and β-glucan peptide (1 mg/ml) were present in the cell culture media throughout the entire 8 h incubation period. To ensure efficient priming by LPS in the absence of serum, 2 μg/ml of recombinant murine LBP was added to LPS stimulated cells. To end the reaction 250 μL of 5X SDS loading buffer was added to each well to capture proteins present in both the cell lysates and supernatants. Samples were homogenized by passing them through a 26-gauge needle, split equally into two tubes and 75 μL of TCEP or H_2_O was added to generate non-reducing and reducing immunoblot samples. After heating to 65 °C for 10 min, proteins were separated by SDS-PAGE on a 4%–12% acrylamide gradient gel (Thermo Fisher) and transgene was detected via immunoblot using a tag-RFP-specific antibody. Since the GSDMD pore is resistant to SDS under non-reducing conditions, oligomerization of NT-GSDMD was indicated by a gel shift toward higher molecular weight.

### Dox-induction of NT-GSDMD

0.5 - 1×10^5^ iBMDMs expressing NT-GSDMD under a Dox-inducible promoter (WT control cells transduced with empty vector or electroporated with non-target sgRNA, RagA KO, caspase-8 KO or RIPK1 KO) were seeded in duplicate or triplicate wells in a black 96-well plate in 200 μl of cDMEM. To induce the transgene, media was exchanged for 200 μl fresh cDMEM containing Dox (0.5 - 2 μg/ml as indicated in Figure legends) and plasma membrane perforation/cell lysis were quantified by PI staining and LDH assay after 8 h or 16 h as described below. For inhibitor studies, cells were pretreated with zVAD (20 μM or 50 μM), RIPK1 inhibitor (1 μM or 5 μM), or RIPK3 inhibitor (1 μM or 5 μM) for 1 h and inhibitors were co-administered during Dox treatment. To induce ROS, cells were co-treated with ROS-inducing agents or PAMPs. 10 μl of cDMEM containing rotenone (5 μM final concentration), TTFA (100 μM final concentration) or sodium arsenite (20 μM final concentration) was added to the cell culture media after 4 h. LPS and β-glucan peptide were present in the cell culture media during the entire stimulation. For direct ROS stimulation, 10 μl of stabilized hydrogen peroxide was added to the cells after 6 h to a final concentration of 625 μM.

To assess fraction of cells expressing NT-GSDMD-tagBFP transgene, cells were detached using 150 μl of PBS + 4 mM EDTA and analyzed via flow cytometry on a BD LSRFortessa Cell Analyzer (BD Biosciences). Transgene expression was detected in the PacBlue channel.

### Induction of caspase-8 and RIPK1-mediatd pyroptosis by TAK1 inhibitor treatment

Co-treatment of cells with TAK1 inhibitor (5z7) and LPS was used to induce caspase-8- and RIPK1- dependent pyroptosis. 1×10^5^ iBMDMs were seeded in duplicate wells in a black 96-well plate in 200 μl of cDMEM. After incubation at 37 °C and 5% CO_2_ for 1-2 h to facilitate cell attachment, media was exchanged for 200 μl fresh cDMEM containing 5z7 (250 nM) and LPS (1 μg/ml). Cells were incubated for another 3 or 6 h at 37 °C and 5% CO_2_ before quantifying cell death by PI staining and LDH release assay.

### Propidium iodide staining and LDH assay

To assess plasma membrane perforation or cellular lysis as a proxy of cell death, we quantified fluorescence after PI staining or lactate dehydrogenase release using the CyQuant LDH cytotoxicity assay kit from Thermo Fisher. Briefly, 10 μl of pre-diluted PI solution was added to the cells (1:300 final dilution) 30 min prior to the end of the stimulation and following a centrifugation step to ensure all cells are in the bottom plane of the wells, (5 min at 400 x g) fluorescence was measured on a Tecan Spark or Biotek Synergy Mx device at an excitation wavelength of 530 nm and an emission wavelength of 617 nm. For the LDH assay, 50 μl of cell culture supernatant per well was transferred into a fresh 96-well plate before adding 50 μl of LDH assay buffer and incubation for 10-20 min at 37 °C. Reaction was stopped by adding 50 μl of LDH stop solution to each well. Absorbance at 490 nm and 680 nm was meaured on a Tecan Spark or Biotek Synergy Mx plate reader. In both assays, signal was normalized to lysis control wells (equal number of cells lysed with detergent-containing lysis buffer).

### Pulldown of oxidized GSDMD

Determination of relative cysteine oxidation was performed as described before with modifications (Saurin et al., 2004). 10×10^6^ iBMDMs expressing NT-GSDMD under a Dox-inducible promoter (empty vector-transduced or RagA KO) were seeded in tissue culture-treated 10 cm dishes in 10 ml of cDMEM and incubated at 37 °C and 5% CO_2_ overnight. Media was exchanged for 10 mL of cDMEM containing 2 μg/ml of Dox to induce the expression of NT-GSDMD for 8 h at 37 °C and 5% CO_2_. As indicated, cells were co-treated with ROS-inducing agents. 500 μl of cDMEM containing rotenone (5 μM final concentration), TTFA (100 μM final concentration) or sodium arsenite (20 μM final concentration) was added to the cell culture media after 4 h. Cells were lifted using PBS + EDTA, washed once in cold PBS, and directly resuspended with lysis buffer (RIPA) containing protease inhibitors and 100 mM N-ethylmaleimide to directly label sulfenic acid-containing proteins at the time of lysis. After incubation on ice for 1 h, lysate was clarified by centrifugation at 16,000 x g at 4°C for 15 min. Protein concentrations were normalized by BCA Assay (Pierce ThermoFisher, Rockford, lL, USA) and excess N-ethylmaleimide was removed by filtration with Zeba spin columns with a 7kDa cut-off (Thermo Fisher). Sulfenic acid residues were selectively reduced with 200 mM sodium arsenite and newly released free thiols were labeled by adding 200 mM 1mM biotin–maleimide. The mixture was incubated at 37°C for one hour. Excess biotin–maleimide was removed using Zeba spin desalting columns. The protein mixture pre-cleared with Sepharose 6B beads (MilliporeSigma) on a rotator at room temperature for two hours. The eluate was further incubated with NeutrAvidin agarose beads (Thermo Scientific Pierce) overnight 4°C to pull down biotin-labeled proteins. NeutrAvidin agarose was washed extensively with wash buffer (50 mM Tris, 600 mM NaCl, 1 mM EDTA, 0.5% NP-40, pH 7.5) and then cold PBS. Enriched proteins were eluted by boiling the NeutrAvidin agarose resin in 4x SDS buffer. Proteins were separated by SDS gel electrophoresis and amount of oxidized NT-GSDMD was assessed by immunoblot using a tagBFP or GSDMD-specific antibody.

### Fluorescence recovery after photobleaching (FRAP)

FRAP experiments were performed on a Zeiss 880 laser scanning confocal microscope using the in-built FRAP module within the microscope control software Zen Black. 0.5 × 10^6^ iBMDMs expressing BFP-tagged NT-GSDMD under the control of a Dox-inducible promoter were plated on a 35 mm μ-Dish (Ibidi; Munich, Germany) in 1 ml of cDMEM and incubated overnight at 37 °C, 5% CO_2_. Media was exchanged for 1 ml of Opti-MEM containing 2 μg/ml Dox for 8 h to induce transgene expression. As indicated, cells were cotreated with ROS-inducing agents or PAMPs. 50 μl of Opti-MEM containing rotenone (5 μM final concentration), TTFA (100 μM final concentration) or sodium arsenite (20 μM final concentration) was added to the cell culture media after 4 h. Following treatments cells were scanned using a 63X oil immersion lens with the 405 nm laser. Regions of interest (ROI) were created at cell membranes and fluorescence bleached by rapid scanning of increased laser power (5-10%) to a bleach depth of 40-60%. Time-lapse images were acquired over a 3-minute time course post-bleaching at 2-second intervals. Images were processed in Zen and FRAP data were fit to a single exponential model using GraphPad Prism.

Data analysis was performed using previously published methods (Haggie and Verkman, 2008). Fluorescence intensities of regions of interest (ROI) in the bleaching area (ROIb = bleached area) were recorded for each time point. The final data was normalized to pre-bleached intensities of the ROIs data and fitted to a single exponential recovery curve. Percent fluorescence recovery (mobile fraction) was calculated from the plateau (Vmax) of the fitted curves normalized to the total bleached fluorescence.

### ROS measurements

0.5 × 10^6^ iBMDMs expressing BFP-tagged NT-GSDMD under the control of a Dox-inducible promoter (RagA KO or WT control cells) were plated on a 35 mm μ-Dish (Ibidi; Munich, Germany) in 1 ml of cDMEM and incubated overnight at 37 °C, 5% CO_2_. Media was exchanged for 1 ml of Opti-MEM containing 2 μg/ml Dox for 8 h to induce transgene expression. As indicated, cells were cotreated with ROS-inducing agents or PAMPs. 50 μl of Opti-MEM containing rotenone (5 μM final concentration), TTFA (100 μM final concentration) or sodium arsenite (20 μM final concentration) was added to the cell culture media after 4 h. LPS (1 μg/ml) and β-glucan peptide (1 mg/ml) were present in the cell culture media throughout the entire 8 h incubation period. To ensure efficient priming by LPS in the absence of serum, 2 μg/ml of recombinant murine LBP was added to LPS stimulated cells. Post-stimulation, cells were loaded with 5μM CM-H_2_DCFDA (Thermofisher Scientific, MA, USA) for 10 mins at 37 °C. Cells were washed three times with PBS and immediately imaged at 37°C on a Zeiss 880 laser scanning confocal microscope using 63X oil immersion lens and 488 nm laser excitation. 5×5 tiled 2D images were obtained from at least three different areas per dish. Images were analyzed and mean fluorescence intensity (MFI) of segmented cells was measured using the intensity analysis function in Image J.

## Acknowledgments

We thank all members of the Kagan lab for helpful discussions. P.D. was supported by a scholarship by the Boehringer Ingelheim Fonds. C.L.E. was supported by a Ragon Early Independence Fellowship. I.H.-B. and E.B. would like to acknowledge funding by the Slovenian Research Agency (ARRS grants J3- 1746 and P4-0176 and young researcher grant). J.R.T. is supported by grants R03DK125630 (NIDDK) and R35GM142683 (NIGMS) and P30DK034854 (NIDDK).

## Supplementary Figure legends

**Supplementary Figure 1.**
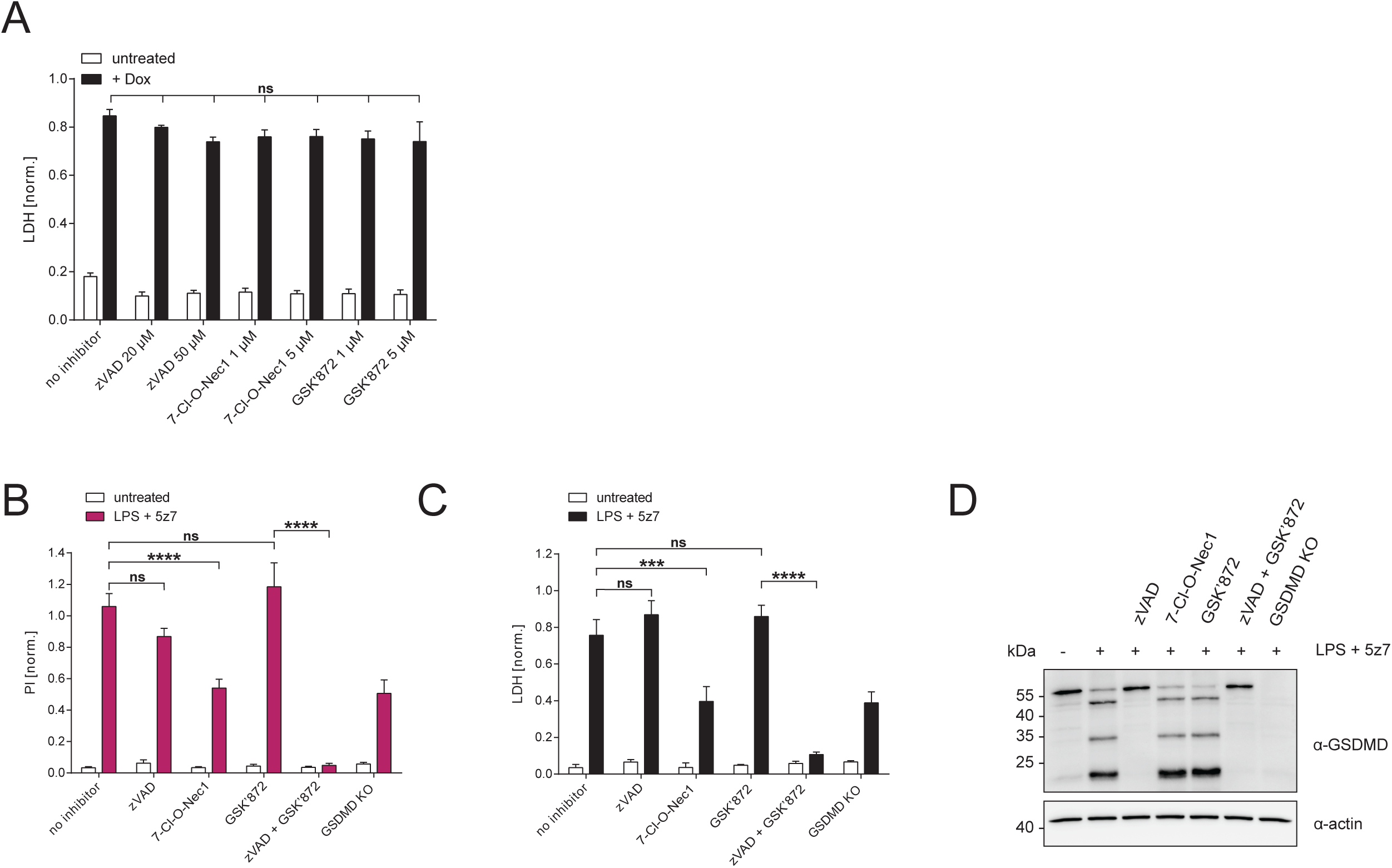
Inhibitors of RIPK1 or caspase-8 do not affect Dox-induced NT-GSDMD-dependent pyroptosis. (A) iBMDMs expressing NT-GSDMD under Dox-inducible promoter were pretreated with pan-caspase inhibitor (zVAD), RIPK1 inhibitor (7-Cl-O-Nec-1) or RIPK3 inhibitor (GSK’872) at indicated concentrations for 1 h. Transgene expression was induced with Dox (0.5 μg/ml) in the presence of inhibitors for 16 h and LDH release into supernatant was quantified. (B,C,D) WT or GSDMD-deficient iBMDMs were pretreated with 50 μM pan-caspase inhibitor (zVAD), 5 μM RIPK1 inhibitor (7-Cl-O-Nec-1) or 5 μM RIPK3 inhibitor (GSK’872) at indicated concentrations for 1 h. Cells were then either left untreated or co-treated with LPS (1 μg/ml) and the TAK1 inhibitor 5z7 (250 nM) for 3 h in the presence of respective inhibitors. PI uptake and LDH release into cell culture supernatants were quantified to assess plasma membrane disruption and cellular lysis, respectively, and cleavage of GSDMD was determined by immunoblot. Data are represented as mean ± SEM of three independent experiments. Immunoblots show representative result of three independent repeats. Statistical significance was determined by two-way ANOVA: *p < 0.05; **p < 0.01; ***p < 0.001; ****p < 0.0001.

## Notes

### Competing Interest Statement

J.C.K. consults for IFM Therapeutics and consults and holds equity in Corner Therapeutics, Larkspur Biosciences and Neumora Therapeutics. None of these relationships influenced the work performed in this study. The other authors declare that they have no competing interests.

